# A frameshift mutation drives divergent biosynthesis of metallophores in *Methylobacterium extorquens*

**DOI:** 10.64898/2026.07.08.737268

**Authors:** Alexa M. Zytnick, Marquis T. Yazzie, Tashi C.E. Liebergesell, Elan H. Tran, Zachary L. Reitz, Aaron W. Puri, Allegra T. Aron, Norma Cecilia Martinez-Gomez

## Abstract

Iron is widely considered the first metallocofactor, evolving as iron-sulfur clusters in early life. While iron-chelating siderophores have been widely characterized across microbial life, the lanthanide-chelating metallophore, methylolanthanin, has only recently been described in *Methylobacterium extorquens* AM1. Methylolanthanin shares structural similarities to the siderophore rhodopetrobactin but contains 4-hydroxybenzoate chelating moieties in place of canonical 3,4-dihydroxybenzoates. Here we compare *M. extorquens* AM1, which produces methylolanthanin, and the closely related *M. extorquens* PA1, which produces rhodopetrobactin. We present a pathway for the biosynthesis of both metallophores and describe the unusual synthesis of methylolanthanin’s 4-HB moieties from tyrosine. We uncover a frameshift mutation in the predicted 3-dehydroshikimate dehydratase, *mllF*, that prevents production of rhodopetrobactin in AM1 through truncation of the catalytically essential N-terminus. We find that deletion of the uncharacterized gene *mllG* reveals a cryptic branch of the pathway, leading to production of both methylolanthanin and rhodopetrobactin. Finally, we discover that rhodopetrobactin production in this mutant is enabled through the activity of a 3-dehydroshikimate dehydratase in a separate biosynthetic gene cluster. These insights highlight an evolutionary mechanism for metallophore diversification through pseudogenization and regulation of distinct biosynthetic gene clusters with shared aromatic intermediates.

## Introduction

Metals are essential for a wide range of biological processes, from acting as signals in regulatory cascades to stabilizing DNA structure.^1–3^ Metals also play key catalytic and structural roles within the majority of enzymes.^3^ While the biological importance of transition metals has been extensively studied, the *f*-block lanthanides have only recently been discovered as biologically relevant metals. XoxF, a methanol dehydrogenase from methylotrophic bacteria, was the first enzyme to be classified as lanthanide-dependent. Since XoxF’s discovery, the field of lanthanide biology has been steadily growing, with additional lanthanide-dependent alcohol dehydrogenases from both methylotrophic and non-methylotrophic organisms being discovered.^4–6^

One way metals such as lanthanides are acquired is through the secretion of small molecule metallophores. This is a well-established strategy across various microorganisms, with iron-chelating siderophores and copper-chelating chalkophores playing essential ecological roles in host-pathogen and plant environments.^7–9^

These secondary metabolites are built by elegant biosynthetic complexes, such as non-ribosomal peptide synthetases (NRPS) or NRPS-polyketide synthases (PKS) hybrids. Some microbial siderophores have been characterized to be NRPS independent and are instead derived from dicarboxylic acid and diamine or amino alcohol units.^10^

Recently, it was discovered that methylolanthanin (MLL) is a lanthanide chelator produced by the model methylotroph *Methylobacterium extorquens* AM1 via an NRPS-independent pathway in response to the limitation of bioavailable lanthanides (Fig. 1A).^11^ MLL is a citrate based metallophore, with 4-hydroxybenzoate (4-HB) moieties and acetylated homospermidine arms. MLL’s unique 4-HB moiety precludes the molecule’s classification into the traditional chemical groups of metallophores: catecholates, phenolates, hydroxamates, carboxylates, or diazeniumdiolates. This moiety is also the key structural difference between methylolanthanin and the catecholate siderophore rhodopetrobactin (RPB) (Fig. 1B).^12^ Interestingly, MLL is not a biological intermediate of RPB, despite its production in response to iron limitation, as the catechol groups of *Bacillus anthracis* Sterne’s petrobactin originate from the direct dehydration of 3-dehydroshikimate to 3,4-dihydroxybenzoate.^13^

**Figure 1.**
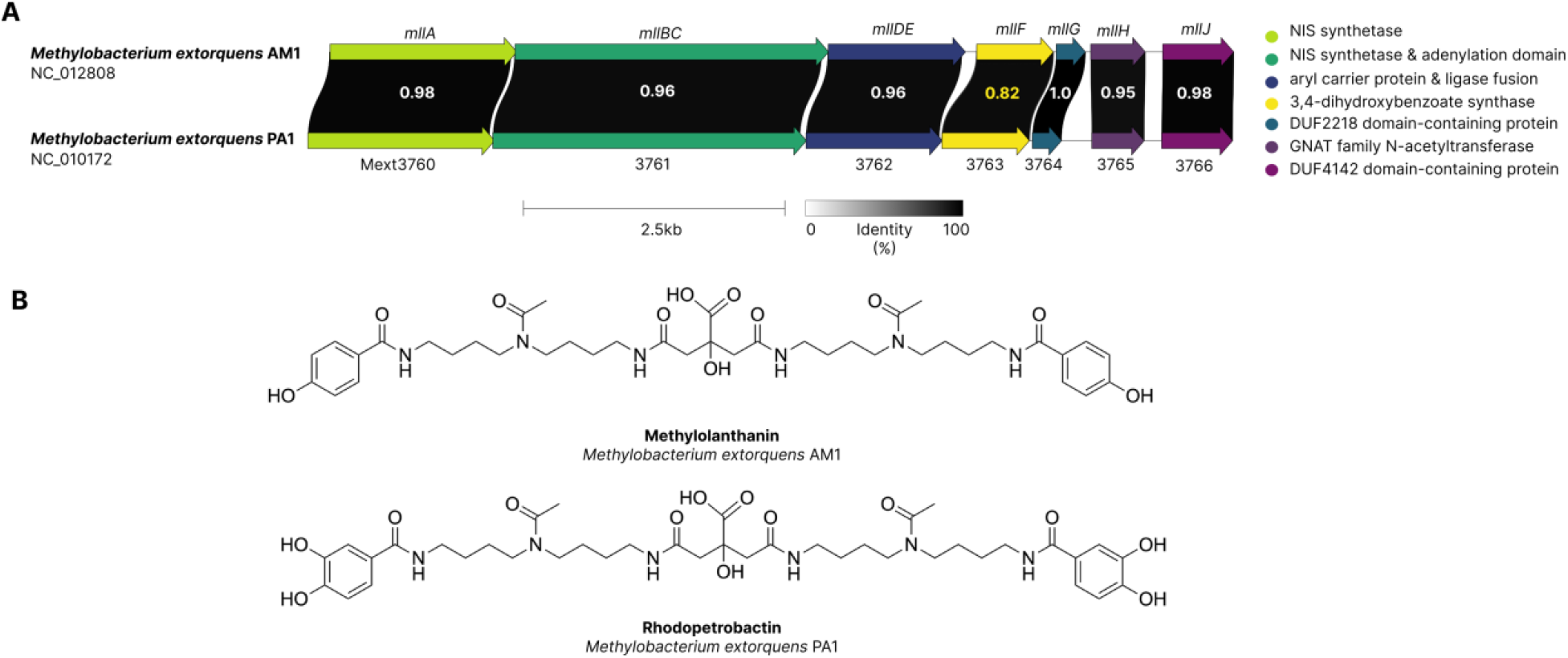
Metallophore loci in *Methylobacterium extorquens* AM1 and PA1. (A) Comparison of the synteny and average nucleotide identities of the biosynthetic gene clusters of methylolanthanin from AM1 and rhodopetrobactin from PA1. The gene with the lowest ANI, *mllF*, is highlighted in yellow. (B) Chemical structures of methylolanthanin and rhodopetrobactin, which differ by the presence of 4-HB vs. 3,4-DHB, respectively.

*Methylobacterium extorquens* PA1 robustly produces RPB (Fig. 1A, B).^14^ We take advantage of this finding as well as the fact that *M. extorquens* AM1 does not produce RPB to compare the biosynthetic routes to these two metallophores. Individual *mll* knockouts in AM1 revealed essential, nonessential, and putative regulatory proteins in the production of MLL. From these data, we present a proposal for the biosynthesis of MLL and RPB, with the aromatic precursor origin controlling the output of the *mll* BGC. From inverse stable isotopic labeling (InverSIL) experiments, we present the biosynthesis of these aromatic precursors, describing the unique production of methylolanthanin’s 4-HB from tyrosine. Through multiple sequence alignments, protein structure prediction, and the production of genetic mutants, we show the 3-dehdryoshikimate dehydratase in the *mll* cluster is dispensable for MLL and RPB biosynthesis, suggesting that pseudogenization of this biosynthetic enzyme allowed for metallophore diversification. Finally, we characterize a mutant that produces both MLL and RPB through utilizing an active 3-dehydroshikimate dehydratase from a different locus. Together, these findings reveal the mechanistic basis of divergent metallophore production from very similar biosynthetic gene clusters.

## Results

### Mutagenesis reveals the function of biosynthetic genes essential for MLL production

To investigate the biosynthesis of MLL, individual deletions of each biosynthetic gene (*mllA, mllBC, mllDE, mllF, mllG, mllH*, and *mllJ*) were made in a Δ*mxaF* background. Mutants were grown in minimal media with methanol as a carbon substrate and supplemented with NdCl_3_ or Nd_2_O_3_. Supernatant extracts were analyzed via non-targeted metabolomics using ultra-high-performance liquid chromatography–tandem mass spectrometry (UHPLC– MS/MS).

*mllF, mllG*, and *mllJ* were found to be dispensable for MLL production (Fig. 2A), while *mllA, mllBC*, and *mllDE* were found to be essential for MLL production (Fig. 2B), consistent with biosynthetic analysis of the petrobactin pathway from *Bacillus anthracis* Sterne.^15,16^ Using petrobactin biosynthesis as a model, we hypothesized MllA to catalyze the addition of 4-HB-homospermidine (**1**) to citrate to form 4-HB-homospermidine-citrate (**2**). MllBC is hypothesized to catalyze the condensation of homospermidine or another 4-HB-homospermidine to 4-HB-homospermidine-citrate to produce the symmetric intermediate di-N-citrylhomospermidine (**3**) or the asymmetric intermediate 4-HB-homospermidine-citrate (**4**) (MllB) coupled to the transfer of 4-HB to a phosphopantetheine group attached to MllD (MllC). MllDE is hypothesized to condense 4-HB-loaded MllD with its substrates (**3**) or (**4**). Finally, MllH is hypothesized to transfer acetyl groups onto amines to produce the final metallophore.

**Figure 2.**
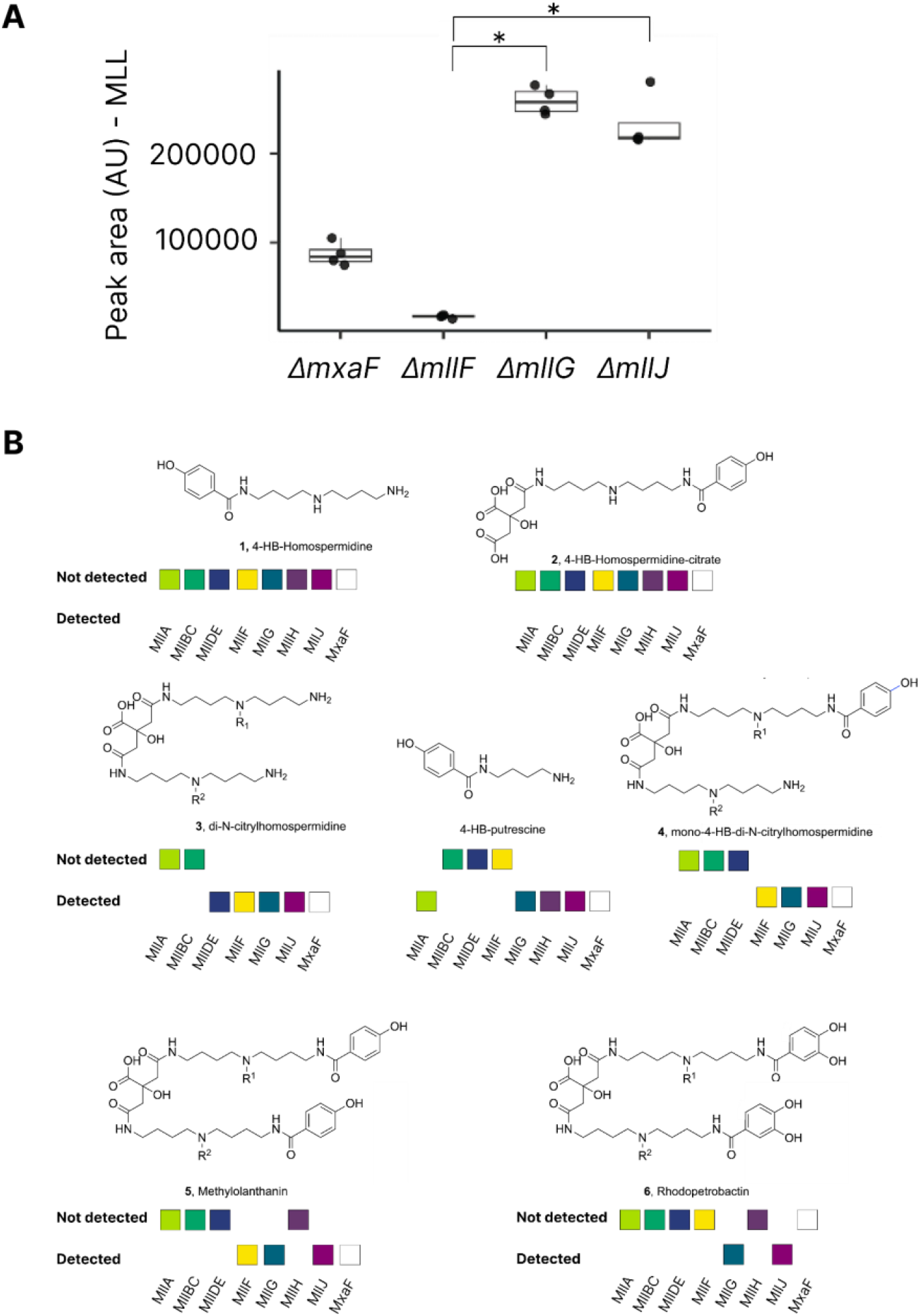
Production of MLL and RPB and their corresponding intermediates in individual *mll* cluster gene knockouts in AM1. (A) Relative abundance of MLL in Δ*mxaF*, Δ*mllF*, Δ*mllG*, and Δ*mllJ* deletion strains. Bars represent the mean of four replicates and error bars represent SD. All growth was carried out in the presence of NdCl_3_. (B) Detection via HPLC-MS/MS of MLL, RPB and proposed MLL intermediates in knockouts of essential *mll* genes compared to *mxaF* control. Peak areas of intermediates can be found in Fig. S4-S12. Growth was carried out in the presence of NdCl_3_ or Nd_2_O_3_.

Interestingly, 4-HB-putrescene was one of the most differentially abundant features between the *mllA* and the *mllBC* deletions (Fig. S1A). While not the hypothesized 4-HB-homospermidine, the absence of 4-HB-putrescene in Δ*mllBC* and Δ*mllDE* is consistent with the proposed functions of MllBCDE in activating and condensing 4-HB. Interestingly, MllF was found to be required for 4-HB-putrescene production (Fig. 2B and Fig. S1B). Additionally, it has been reported that 4-HB-homospermidine is difficult to detect via MS.^16^ We also do not detect 4-HB-homospermidine-citrate (**2**) in any samples because MllA is predicated to catalyze this reaction and the MllBC and MllDE fusions are necessary to produce the 4-HB-homospermidine precursor of (**2**).

Intermediate (**3**) was not found in Δ*mllA* and Δ*mllBC* but was found in Δ*mllDE* (Fig. 2B), which highlights the role of *mllDE* in 4-HB ligation. Deletion of *mllH* resulted in intermediates lacking acetyl groups, confirming its proposed role as an N-acetyltransferase (Fi. 2B and Fig. S9). Due to their co-upregulation with the *mll* cluster, META1p5050 and META1p5051 were identified as potential hydroxylases and decarboxylases acting upon the aromatic precursor of 4-HB, but deletion of either gene still yielded MLL production (Fig. S2).

### Inverse labeling towards elucidating the origin of 4-hydroxybenzoate

As the key difference between MLL and RPB is the replacement of two 3,4-DHB moieties with 4-HB moieties, the origin of 4-HB was investigated through utilizing InverSIL.^17^ The Δ*mllF* mutant, which produces only MLL (Fig. 2A), was grown in minimal media with ^13^C-substituted methanol as a carbon source and supplemented with unlabeled putative aromatic intermediates to detect for incorporation via LC-HRMS/MS. Interestingly, when precursors that are eventually incorporated are added, MLL is produced in much greater amounts than the wildtype, suggesting aromatic precursors are in limited supply.

AM1 does not contain a UbiC homolog, which converts chorismate to 4-HB, and would be the assumed source of this aromatic precursor for MLL. A novel pathway for the biosynthesis of 4-HB in *Klebsiella oxytoca* was characterized to originate from tyrosine, proceeding through 4-hydroxyphenylpyruvate, 4-hydroxyphenyllactate, 4-hydroxymandelate, 4-hydroxybenzaldehyde, with a final oxidation to 4-HB.^18^ Through inverse labeling we see incorporation of each of these intermediates from this pathway into MLL, from tyrosine to 4-HB, apart from 4-hydroxyphenyllactate (Fig. 3, Table S1).

**Figure 3.**
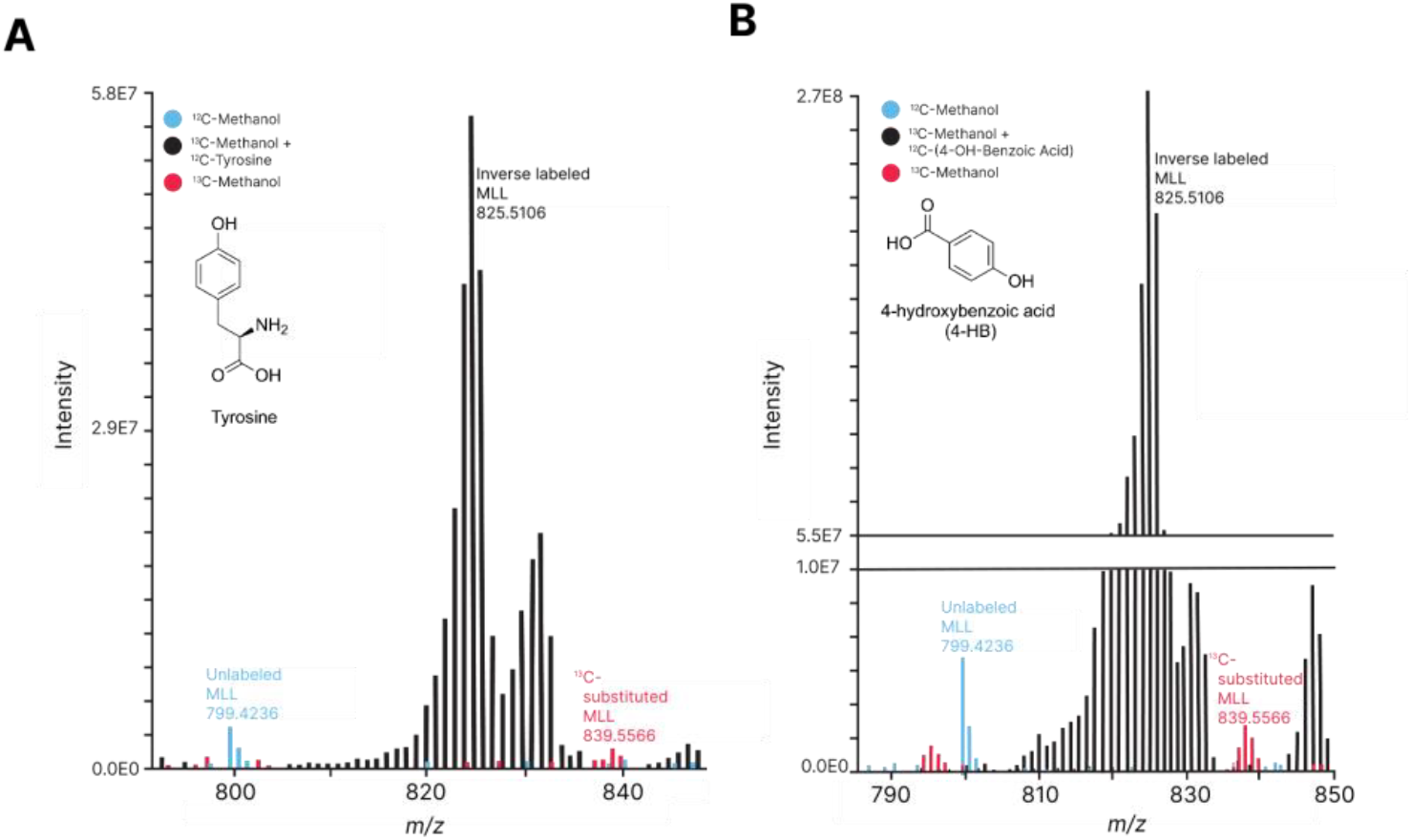
Incorporation of aromatic intermediates into MLL in a *mllF* deletion strain. (A) Incorporation of tyrosine into MLL. (B) Incorporation of 4-HB into MLL. Additional intermediates tested can be found in Table S1.

### *Nonessential* mll *genes control alternative aromatic precursor routing and regulation*

The production of methylolanthanin by *mllF, mllG*, and *mllJ* deletion mutants (Fig. 2A) led us to investigate the roles of their protein products. MllF is annotated as a sugar phosphate isomerase/epimerase family protein and shares similarity with the 3-dehydroshikimate dehydratase AsbF from petrobactin producer *Bacillus anthracis* Sterne.^15^ A blastn search of *mllF* against RefSeq genomes showed nucleotide sequence similarity to other AsbF homologs across methylotrophs, including PA1. A multiple sequence alignment of AsbF homologs from AM1, PA1, and *Methylorubrum aminovorans* NBRC 15686 revealed frameshift mutations in AM1 and *M. aminovorans* NBRC 15686 that are absent in PA1 and other methylotrophs (Fig. 4A). These frameshifts truncate the canonical N-terminus of the protein, however, a downstream in-frame start codon is predicted to permit expression of a folded core protein.

**Figure 4.**
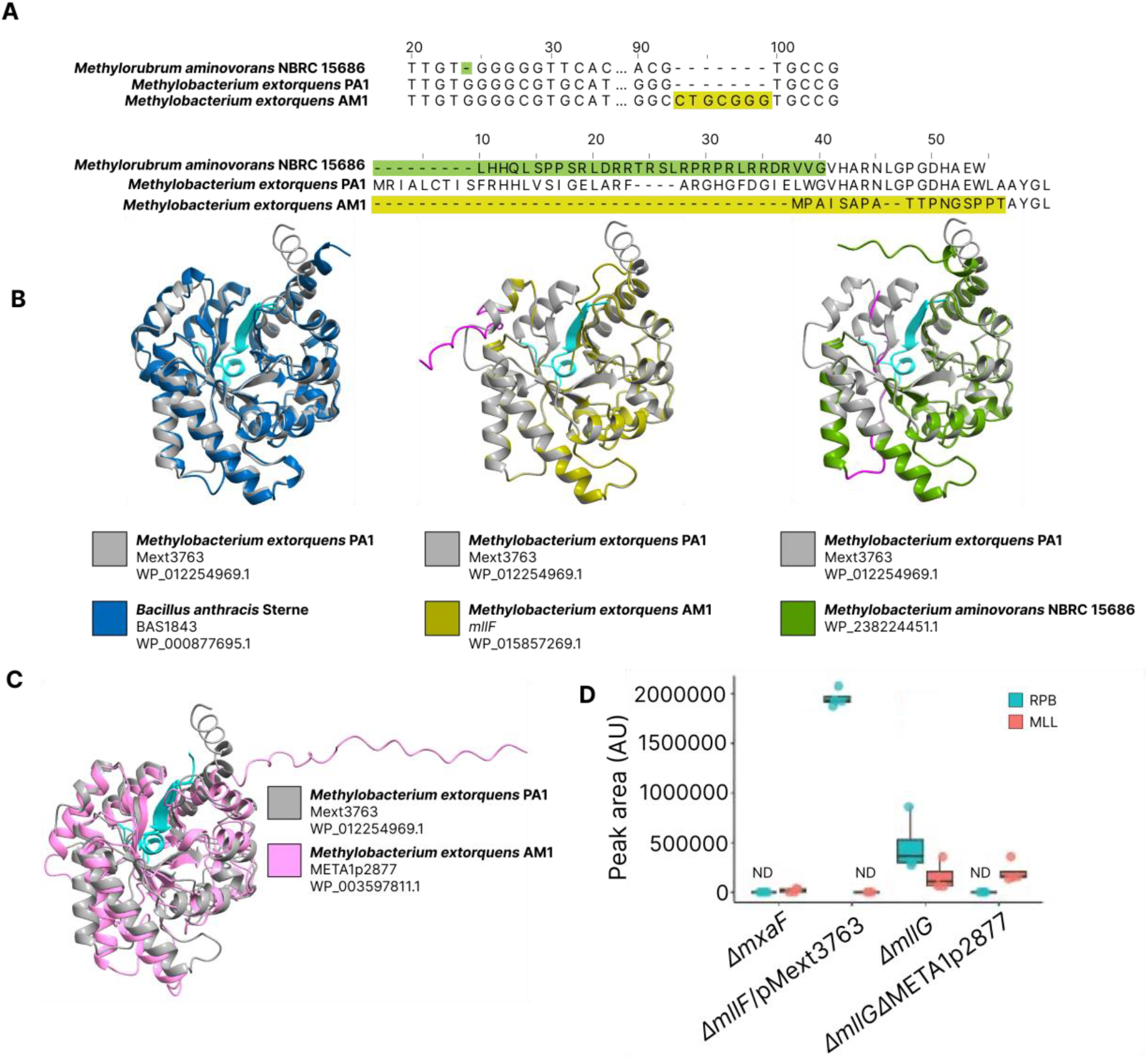
Pseudogenization of the 3-dehydroshikimate dehydratase within the *mll* cluster confers MLL production instead of RPB production. (A) Multiple sequence alignments at the nucleotide and amino acid levels of *mllF* from AM1 and its homologs in PA1 and *M. aminovorans* NBRC 15686. Frameshift mutations, and the resultant truncations in the translated protein, at the N-terminus are highlighted in green and yellow. (B) Overlays of AlphaFold3 models of MllF from AM1, its homologs in PA1 and *M. aminovorans* NBRC 15686, and the crystal structure of AsbF from *B. anthracis* Sterne (3DX5). Correctly folded N-termini are highlighted in cyan and disordered N-termini are highlighted in magenta. (C) Alphafold3 models of a putative 3-dehydroshikimate dehydratase, META1p2877, from AM1 and the MllF homolog from PA1. Correctly folded N-termini are highlighted in cyan. (D) Relative abundance of MLL (salmon boxes) and/or RPB (blue boxes) in Δ*mllF*, Δ*mllF* expressing the PA1 *mllF* homolog Mext3763, Δ*mllG*, and a Δ*mllG*ΔMETA1p2877 double deletion mutant. Bars represent the mean of four replicates and error bars represent SD.

AlphaFold modeling of MllF, homologs from PA1 and *M. aminovorans* NBRC 15686, overlaid with the crystal structure of AsbF from *Bacillus anthracis* Sterne reveal that AsbF and its PA1 homolog share a matching N-terminal beta sheet and loop at the dimer interface (Fig. 4B), interactions that are documented as essential for activity in this family of enzymes.^19^ The homologous proteins in AM1 and *M. aminovorans* NBRC 15686 lack one or both of these N terminal features due to the truncation caused by the aforementioned frameshift mutations (Fig. 4A, B). This leads to catalytically inactive forms of these enzymes, explaining the lack of production of RPB in AM1 and the capacity of the AM1 *mllF* deletion mutant to still produce MLL (Fig. 2A). We verify that the misfolded N-terminus of *mllF* is the cause of MLL production through heterologously expressing the PA1 *mllF* homolog in a Δ*mllF* background in AM1 (Fig. 2D). We find that expression of the active enzyme from PA1, Mext3763, leads to the production of RPB instead of MLL (Fig. 4D). While *M. extorquens* AM1 does not naturally produce RPB under the tested conditions, deletion of *mllG* when grown under Nd_2_O_3_ led to production of RPB in addition to the production of MLL (Fig. 2B and Fig. S5). We find that production of RPB is conferred through the activity of a putative 3-dehydroshikimate dehydratase, META1p2877, that is located in a locus outside of the *mll* cluster. META1p2877 is predicted to retain a folded N-terminus and therefore remain active (Fig. 4C). Deletion of META1p2877 in a *mllG* deletion background abolished the single mutant’s capacity to produce RPB, denoting that META1p2877 is responsible for the production of the 3,4-DHB incorporated into RPB produced by the *mllG* single mutant (Fig. 4D). In addition, the deletion of META1p2877 had no significant effect on the relative abundance of MLL (Fig 4D).

## Discussion

Iron is considered one of the oldest metallocofactors.^20-22^ Lanthanide dependent enzymes and systems for lanthanide uptake likely evolved much later than iron enzymes and uptake systems. During the Phanerozoic to modern eras, acidic environments and terrestrial weathering made lanthanides more available, and even produced Fe-oxyhydroxides that acted as sinks for light lanthanides.^23–26^ Here, we present insights into the evolution of lanthanide and iron metallophores in *M. extorquens* AM1 and *M. extorquens* PA1. Simultaneous work from Gutenthaler-Tietze *et al* has corroborated the production of MLL in AM1, RPB in PA1, and the presence of the frameshift in the AsbF homolog MllF.^27^

We find the biosynthesis of the cores of RPB and MLL proceed similarly to that of the siderophore petrobactin, but with homospermidine instead of spermidine. We find MllBC and MllDE are likely necessary to produce 4-HB-homospermidine from 4-HB and homospermidine (Fig. 5). While MllA is predicted to then catalyze the addition of citrate to 4-HB-homospermidine, **2**, this intermediate was not detected. This may be due to the MllBC and MllDE fusions, in contrast to their standalone homologs in the petrobactin biosynthesis pathway (AsbB, AsbC, AsbD, and AsbE).^16^ Interestingly, the RPB biosynthetic gene cluster from *Rhodospeduomonas palustris* TIE-1 exhibits an intermediate fusion phenotype, as it possesses independent AsbB and AsbC homologs but an AsbDE fusion.^12^ Separation of these genes in the *mll* cluster would be necessary to detect **2**, as they are necessary to produce its 4-HB-homospermidine precursor. This suggests that metallophore diversification within the petrobactin-like family may be accompanied by progressive fusion of biosynthetic enzymes.

**Figure 5.**
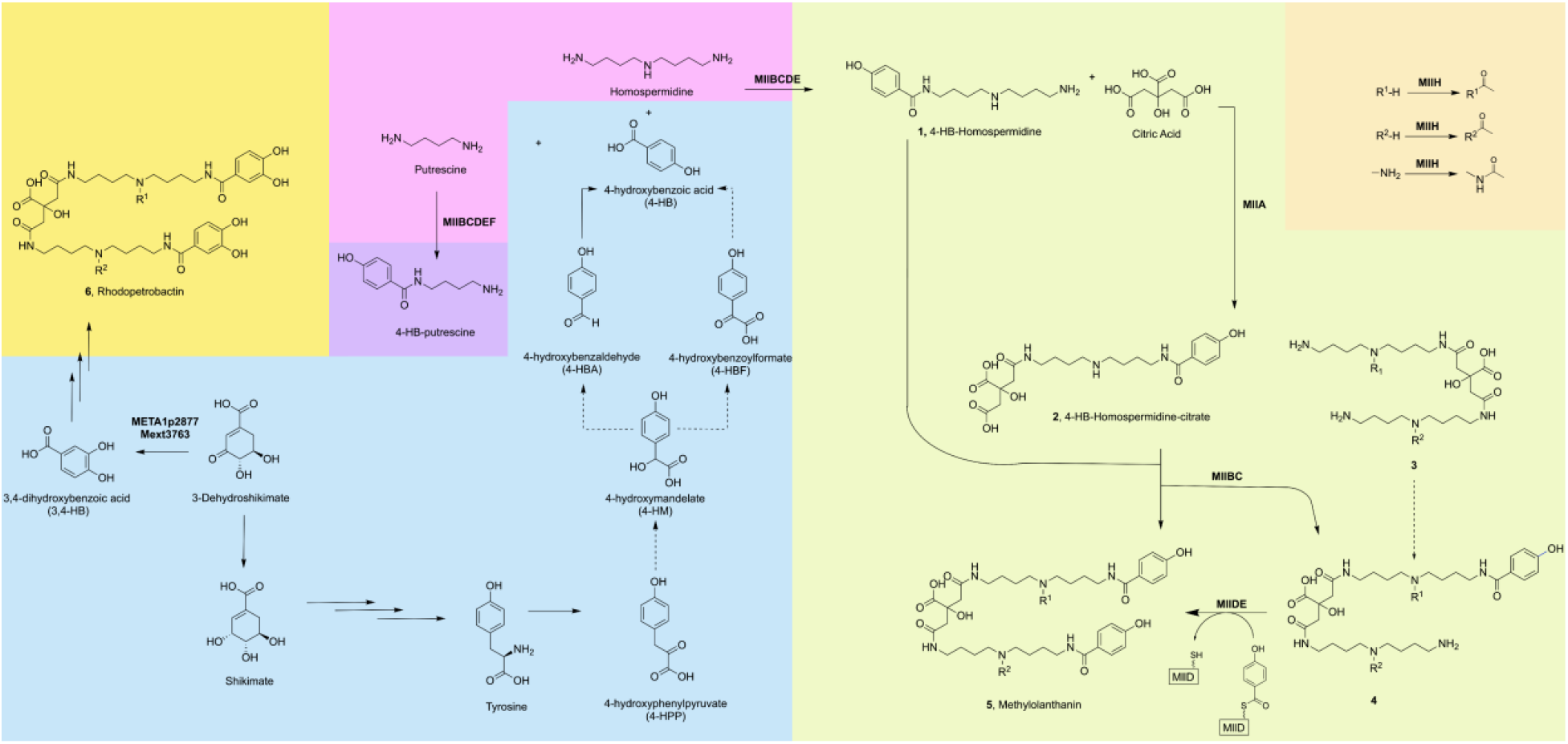
Proposed biosynthetic scheme of MLL, RPB, and 4-HB-putrescine. Formation of the 4-HB and 3,4-DHB chelating moieties (blue), and the role of MllABCDEH in producing the citrate core (green), MLL (green), performing acetylation (orange), and producing 4-HB-putrescine (purple) from polyamine precursors (pink) as determined by InverSIL and HPLC-MS/MS. Dashed arrows represent putative reactions between intermediates.

We find MllBC to then condense another 4-HB-homospermidine to the citrate core, forming MLL, **5**. Alternatively, we find another branch of MLL biosynthesis to be active, in which MllDE catalyzes the addition of a second 4-HB to **4**, an intermediate that is only mono-substituted with 4-HB. We find **4** to originate from dihomospermidine-citrate, **3**, which is congruent with previously described asymmetric petrobactin biosynthesis, although we do not detect the earlier intermediates from this branch of the pathway. We confirm that MllH is responsible for N-acetylation (Fig. S9) and find mono-, di-, and tri-acetylated versions of **4** and **3**.

The AsbF homolog, MllF, and the uncharacterized proteins MllJ and MllG were not found to be essential for MLL production (Fig. 2A). MllF was found to be essential for the production of 4-HB-putrescine (Fig. 2B), which is structurally similar but distinct from the polyamine catechol siderophore aminochelin (2,3-DHB-putrescine).^28^ Dating back to the ancestral stock of *M. extorquens* AM1 from 1960,^29^ *mllF* exhibits a frameshift mutation that renders its protein product inactive (Fig. 4A). Heterologous expression of its functional homolog from PA1 allowed for production of RPB over MLL without restoring 4-HB-putrescine production (Fig. 4D, Fig. S1B). AM1 has seemingly evolved away from the canonical RPB biosynthetic strategy by eliminating the need for AsbF-dependent, 3-dehydroshikimate dehydratase activity. As a consequence, MllF has lost its catalytic function but has been retained because it performs a new role in an MLL-associated aromatic polyamine pathway. Within this framework, MllF does not function as a source of 4-HB but rather as a pseudoenzyme that promotes efficient flux through the MLL-producing branch. Deletion of *mllF* therefore reduces, but does not abolish, MLL production (Fig. 2A) while completely eliminating detectable 4-HB-putrescine (Fig. 2B), a phenotype expected if 4-HB-putrescine is a shunt product or overflow metabolite rather than an obligate precursor. The ability of a catalytically active PA1 MllF homolog to restore RPB production but not 4-HB-putrescine production (Fig. S1) further supports the idea that active AsbF chemistry drives the ancestral RPB pathway, whereas the inactive AM1 MllF has been repurposed for a distinct function associated with 4-HB-polyamines and MLL biosynthesis.

We investigate the origin of MLL’s chelating moieties via InverSIL. We find that 4-HB is incorporated into MLL, is not incorporated into RPB, and that 4-HB originates from tyrosine and not from canonical catechol biosynthesis pathways (Fig. 3). 4-HB is produced from tyrosine via a novel route through 4-hydroxyphenylpyruvate, 4-hydroxymandelate, and 4-hydroxybenzaldehyde (Fig. 5). While no annotated 4-HPP dioxygenase was found in the AM1 genome, it has been reported that 4-HPP can be converted to 4-HBA bioinorganically with iron(IV)-oxido and iron(III)-hydroxido species.^30^ AM1 likely then converts 4-HBA to 4-HB through an unspecific aldehyde dehydrogenase, as subsets of this enzyme family has been shown to be promiscuous towards aromatic substrates.^31^

Lastly, we find that RPB and MLL may be linked evolutionarily through regulation, as DUF genes *mllJ* and *mllG* are not biosynthetically essential for production of MLL and because the *mllG* deletion produces both RPB and MLL (Fig. 2A). Indeed, we find that MllG may control RPB production through recruiting an AsbF homolog in a distant locus, META1p2877, for 3,4-DHB production, as a double deletion of META1p2877 and *mllG* fails to produce RPB (Fig. 4D). As PA1 also has *mllJ* and *mllG* (Fig. 1A), they may serve as regulatory switches to control the flow of aromatic intermediates to either RPB biosynthesis or other pathways. Interestingly MllG shows structural similarity to the uncharacterized protein PA4090 from *Pseudomonas aeruginosa* that was found to be differentially expressed in response to iron and the production of the siderophore pyoverdine.^32^ Coupled with the knowledge that the MLL cluster is responsive to iron concentrations,^11^ MllG may further play a role in regulating the production of RPB over MLL in iron limiting conditions. Future work that evaluates the relationship between these proteins will surely provide more insights into the regulatory network underlying metallophore diversity and biosynthesis.

## Acknowledgements

This material is based upon work supported by the NSF under Grants 2127732 and 2223730 (N.C.M.-G.). A.M.Z. was supported by the U.S. Department of Defense (National Defense Science and Engineering Graduate Fellowship). E.H.T was supported by the NSF (Graduate Research Fellowship Program). A.T.A. and M.T.Y. were supported by NSF CHE-2441115. T.C.E.L. was supported by NIH training grant T32 AI055434. Work in the Puri Lab was supported by Simons Foundation Early Career Investigator in Aquatic Microbial Ecology and Evolution Award 00006628 (to A.W.P.) and NSF CAREER Award 2339190 (to A.W.P.).

## Materials and Methods

### General Working and Culturing Conditions

MilliQ grade water and HPLC grade solvents were used in this work, unless otherwise stated. For mass spectrometry and related experiments, LCMS grade solvents were used. Culturing occurred under sterile conditions. *M. extorquens* AM1 strains [Δ*mxaF*, ΔmxaFΔmll,^11^ Δ*mxaF*Δ*mllA*, Δ*mxaF*Δ*mllBC*, Δ*mxaF*Δ*mllDE*, Δ*mxaF*Δ*mllF*, Δ*mxaF*Δ*mllG*, and Δ*mxaF*Δ*mllH*, Δ*mxaF*Δ*mllJ*, Δ*mxaF*ΔMETA1p2877, Δ*mxaF*/pMext3763, Δ*mxaF*ΔMETA1p5050, and Δ*mxaF*ΔMETA1p5051] were grown in MP medium.^33^ 3 mL cultures with 15 mM succinate (MilliporeSigma) were grown overnight in 15 mL round bottom glass culture tubes (Thermo Fisher Scientific) at 30°C and 200 rpm and then diluted into the desired volume of 50 mL of fresh MP to an OD600 of 0.1. Bulk cultures were grown with 50 mM MeOH as a carbon source and 2 μM NdCl_3_ or Nd_2_O_3_ as lanthanide sources. When necessary, sucrose, and kanamycin were added to final concentrations of 5%, 50 μg/mL, and 50 μg/mL, respectively. The above conditions were used unless otherwise noted.

### DNA Manipulation

All fragments were amplified with either Q5 DNA polymerase or Phusion DNA polymerase (New England Biolabs). The Mext3763 overexpression vector pAZ05 was generated using Q5 polymerase to amplify fragments from PA1 gDNA and the pTE1877 backbone and these fragments were digested and ligated together using restriction enzymes and T4 ligase (New England Biolabs).^34^ Deletion vectors were generated by amplifying a pCM433kanT backbone and ∼500 bp flanking regions upstream and downstream of target genes from AM1 gDNA. Fragments for individual *mll* BGC deletions and the META1p2877 deletion were fused using Gibson Assembly Master Mix (New England Biolabs). Constructs were transformed into electrocompetent DH5α or 10β *E. coli* (New England Biolabs), plated on selective media, and purified using the GeneJET Plasmid Miniprep Kit (Thermo Fischer Scientific). For deletion vectors, counterselection was achieved through patching transformants onto Hypho^33^ succinate sucrose and Hypho succinate kanamycin plates and selecting colonies with a successful double crossover event (growth on sucrose, no growth on kanamycin). Full plasmid sequencing or PCR product sequencing (Plasmidsaurus) confirmed the sequence of all constructs and the deletion of target genes.

### Bioinformatic Analysis of ORFs and Protein Structure

Biosynthetic gene cluster comparisons were constructed using clinker.^35^ A NCBI blastn search against RefSeq genomes was used to compile a list of *mllF* homologs.^36,37^ Alternative ORFs for *mllF* were discovered using NCBI ORFfinder with AM1’s genome (RefSeq accession NC_012808). Protein structures were predicted using the AlphaFold 3 server.^38^ Multiple sequence alignments were performed using Clustal Omega.^39,40^

### Mass spectrometry analysis of AM1 biosynthesis mutants for MLL, RPB and biosynthetic intermediates

Samples were received and processed in three separate batches: batch 1 contained all *mll* mutants (Δ*mllA*, Δ*mllBC*, Δ*mllDE*, Δ*mllF*, Δ*mllG*, Δ*mllH*, Δ*mllJ* and Δ*mxaF*); batch 2 contained Δ*mllF*, Δ*mllG*, Δ*mllJ*, ΔMETA1p5050, and ΔMETA1p5051; batch 3 contained Δ*mllF*, Δ*mllG*, Δ*mllF*/pMext3763, and Δ*mllG*ΔMETA1p2877. Detailed descriptions of each batch are provided on Github. All *M. extorquens* AM1 strains were pelleted at 5,000 x g for ten minutes prior to shipment of supernatant at -80 °C. Once thawed, 25-50 mL of supernatant were subjected to solid phase extraction (SPE) using Chromabond HLB sorbents and a protocol aligned with the package insert. Briefly, HLB columns were conditioned with methanol then equilibrated with 3% MeOH in water prior to loading supernatant onto the column. The column was washed with 3% MeOH in water, dried with vacuum, and then eluted with 80% methanol in water. Samples were concentrated then resuspended in 200 µL of 80% methanol with 1µM sulfadimethoxine as the internal standard (batches 1 and 3). Samples in batch 2 were reconstituted to 1mg/mL based on dry sample biomass. For samples in batches 2 and 3, 1µM sulfamethazine was added as an extraction standard prior to SPE.

2-5 µL of sample was injected into a Vanquish coupled to a Q Exactive HF quadrupole orbitrap mass spectrometer (Thermo Scientific). A Waters Acquity C18 UPLC HSS T3 1.8μm 2.1×100mm was used for chromatography. The mobile phase consisted of solvent A (water +0.1% formic acid (FA)) and solvent B (acetonitrile + 0.1% FA). The flow rate was set to 0.5 mL/min. After injection, the samples were eluted with the following method: 0 to 6 min 2% to 50% B, 6 to 10 min 50 to 99% B, followed by a 3 min washout phase at 99% B and a 3-min re-equilibration phase at 2% B. After 10 minutes from the start of the run, the divert valve switched to divert the LC flow from MS to waste. Data-dependent acquisition (DDA) of MS/MS spectra was performed in positive electrospray ionization mode. Full MS was collected with 120,000 resolution, 1e6 automatic gain control target, maximum injection time 100ms for the 150 to 1500 m/z range. The dd-MS^2^ was collected with 15,000 resolution, 5e5 automatic gain control target, maximum injection time 200ms, TopN 5, isolation window 1.0 m/z, and normalized collision energy stepped 25, 35 and 45. A QC mixture containing sulfamethazine, sulfamethizole, sulfachloropyridazine, sulfadimethoxine, and coumarin-314 was run every ten samples, along with method blanks.

### Feature Finding and Feature Based Molecular Networking

Profile spectra (.raw) were converted to .mzML files using MSconvert. mzmine (4.7.29) was used to detect features across batches 1 & 2, and mzmine (4.9.14) was used for batch 3. All mzmine processing files and parameters can be found in Github. GNPS2 Feature Based Molecular Networking (FBMN) was used for all data visualization and can be accessed at the following links: https://gnps2.org/status?task=4cdb0a26b5d94b258b83629ab2ad9eb6 (batch 1: presence/abscence of MLL, RPB and biosynthetic intermediates across BGC mutants); https://gnps2.org/status?task=673f01deb5c1428588495bba2c630725 (batch 2: relative abundance of MLL across ΔmllF, ΔmllG, ΔmllJ); https://gnps2.org/status?task=8236c5b380d1425293d7517c3d42d93a (batch 3: relative abundance of MLL and RPB across Δ*mllF*, Δ*mllG*, Δ*mllF*/pMext3763, and Δ*mllG*ΔMETA1p2877). Batch 1 (presence/absence of MLL, RPB and biosynthetic intermediates) was blank subtracted and imputed, and normalized, as previously described.^41^ Batch 2 (relative abundance of MLL across Δ*mllF*, Δ*mllG*, Δ*mllJ*, ΔMETA1p5050, and ΔMETA1p5051) was not blank subtracted or imputed.

### Inverse Labeling and Detection of Methylolanthanin and Rhodopetrobactin

Exponentially growing cultures were pelleted at 4,000 rpm for ten minutes and resuspended in fresh ammonium mineral salts (AMS) medium with no carbon source. Modified AMS contains 0.2 g L^-1^ MgSO_4_·7H_2_O, 0.2 g L^-1^ CaCl_2_·6H_2_O, 0.5 g L^-1^ NH_4_Cl, 30 µM LaCl_3_, and 1X trace elements. 500X trace elements contains 1.0 g L^-1^Na_2_-EDTA, 2.0 g L^-1^ FeSO_4_·7H_2_O, 0.8 g L^1^ ZnSO_4_·7H_2_O, 0.03 g L^-1^ MnCl_2_·4H_2_O, 0.03 g L^-1^ H_3_BO_3_, 0.2 g L^-1^CoCl_2_·6H_2_O, 0.6 g L^1^ CuCl_2_·2H_2_O, 0.02 g L^-1^ NiCl_2_·6H_2_O, and 0.05 g L^-1^ Na_2_Mo·2H_2_O. A final concentration of 4 mM phosphate buffer pH 6.8 and 50 mM ^12^C-or ^13^C-methanol were added prior to use. Subsequently, three separate six-milliliter cultures were inoculated with the resuspended strain to a starting OD (600 nm) of 0.05. ^12^C-methanol was added to one culture, ^13^C-methanol was added to another culture, and ^13^C-methanol plus the ^12^C-precursor (final concentration of precursor = 100 µM) was added to the final culture. All cultures were grown at 30C and shaken at 200 rpm. Metabolite extraction and LC-HRMS/MS were performed as previously described.

## Data availability

All mass spectrometry .mzML files and mzmine-processed results are available in the mass spectrometry interactive virtual environment (MassIVE) under massive.ucsd.edu with project identifiers: MSV000102363 (Batch 1: presence/absence of MLL, RPB and biosynthetic intermediates in BGC mutants), MSV000102359 (Batch 3: MLL and RPB relative abundance for mllF, mllG, mllF/pMext3763, and ΔmllGΔMETA1p2877), MSV000102362 (Batch 2: MLL relative abundance for mllF, mllG, mllJ, 5050 and 5051). All relevant code and analysis is found at https://github.com/marquisyazzie/Biosynthesis-of-MLL.

