## Supplementary Information for "A frameshift mutation drives divergent biosynthesis of metallophores in *Methylobacterium extorquens*"

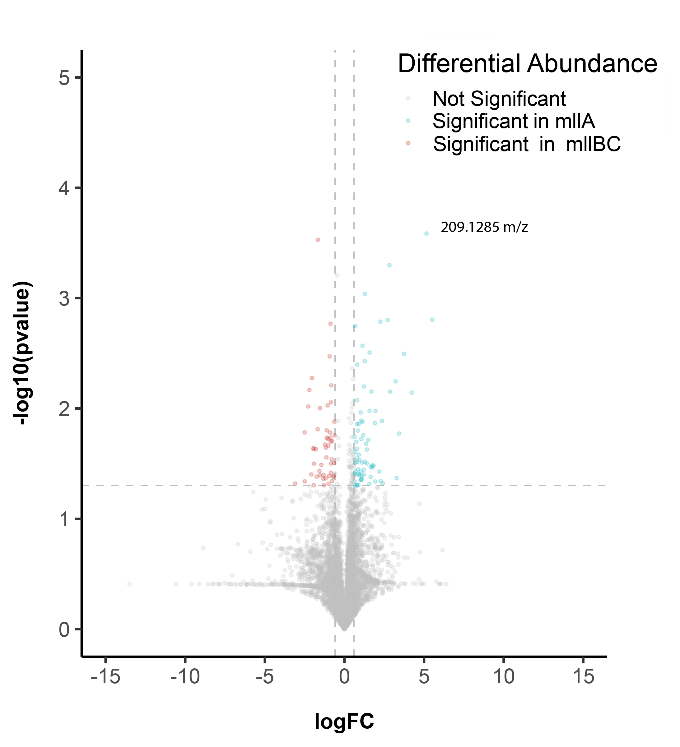

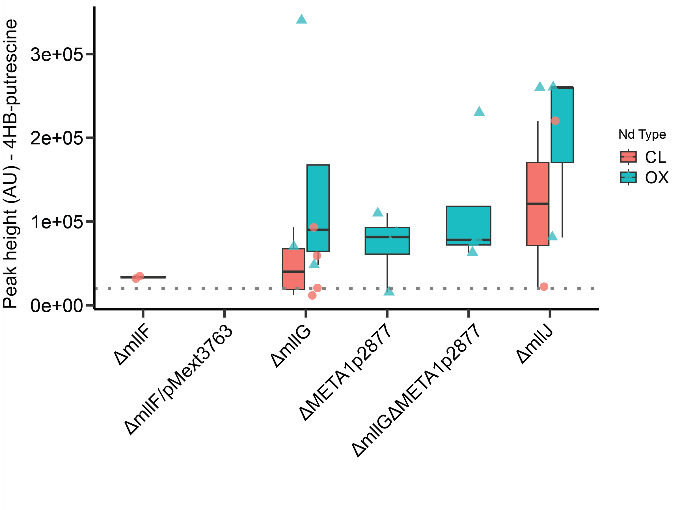

**Figure S1**. (A) Volcano plot of Δ*mllA* vs Δ*mllBC* with 4-HB-putrescine (209.1285 *m/z* observed mass). (B) Relative abundance of 4-HB-putrescine in various *mll* mutants.

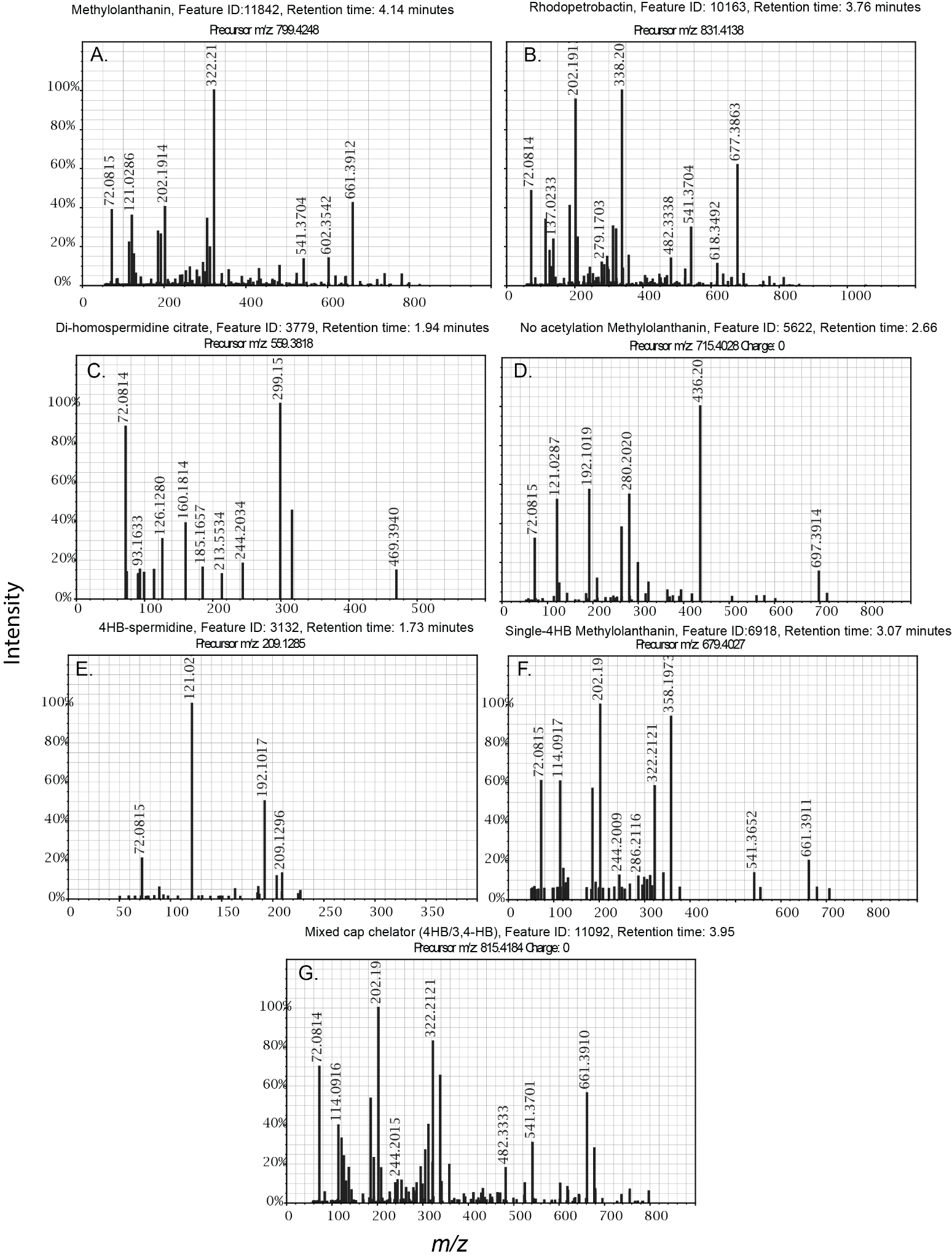

**Figure S2.** (A-G). MS/MS of MLL, RPB, and manually annotated intermediates from scheme

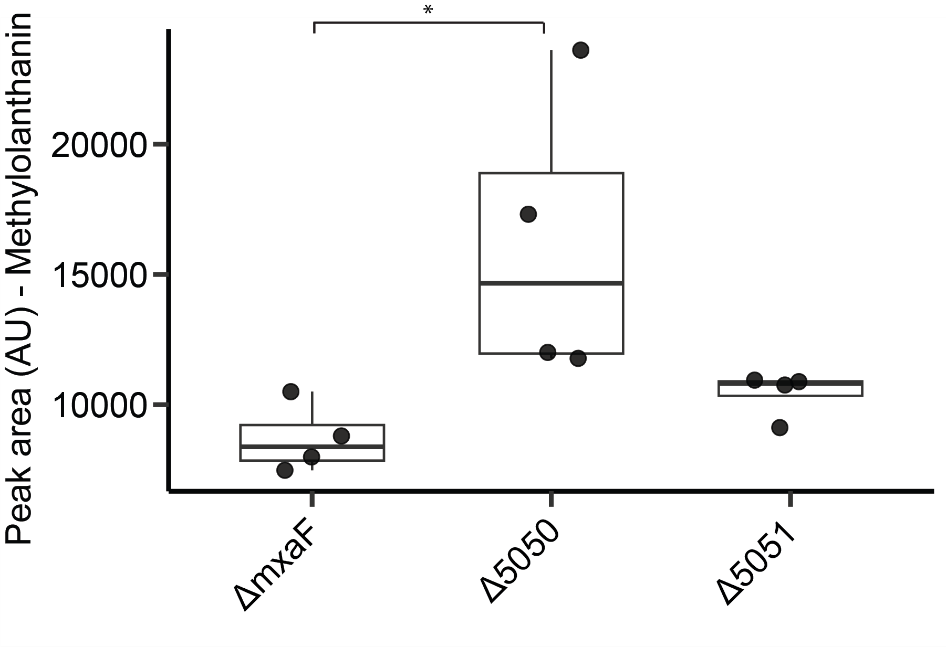

**Figure S3.** ΔMETA1p5050 and Δ5051 still produce MLL at relative abundances comparable to Δ*mxaF.*

**Table S1.** Additional intermediates tested via InverSIL

| **Intermediate tested** | **Incorporation into MLL** |
| --- | --- |
| 4-hydroxyphenylpyruvate | Yes |
| 4-hydroxyphenylacetate | No |
| 4-hydroxymandelate | Yes |
| 4-hydroxybenzaldehyde | Yes |
| 4-coumarate | Yes |
| 4-hydroxybenzoylformate | Yes |
| acetate | Yes |
| shikimate | Yes |
| anthranilate | Yes |
| phenylalanine | No |
| tryptophan | No |
| 2,3-DHB | No |
| 3-dehydroshikimate | No |

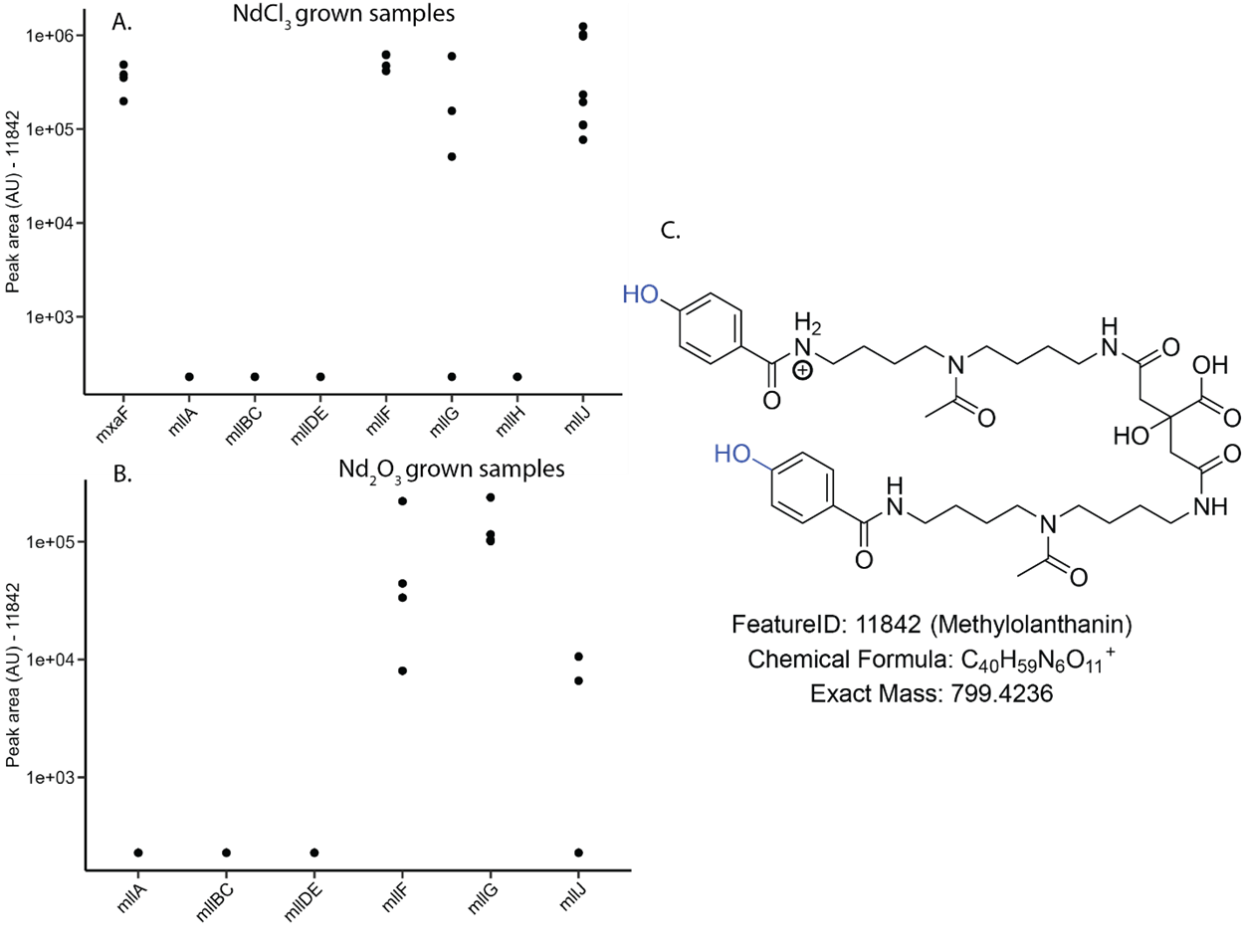

**Figure S4.** Imputed peak areas of MLL across *mll* mutants in (A) NdCl_3_ grown samples and (B) Nd_2_O_3_ grown samples. (C) Structure of MLL (feature 11842). This data was used to determine the presence/absence of this feature in Figure 2B. Peak areas have not been normalized to OD and originate from different sample growths.

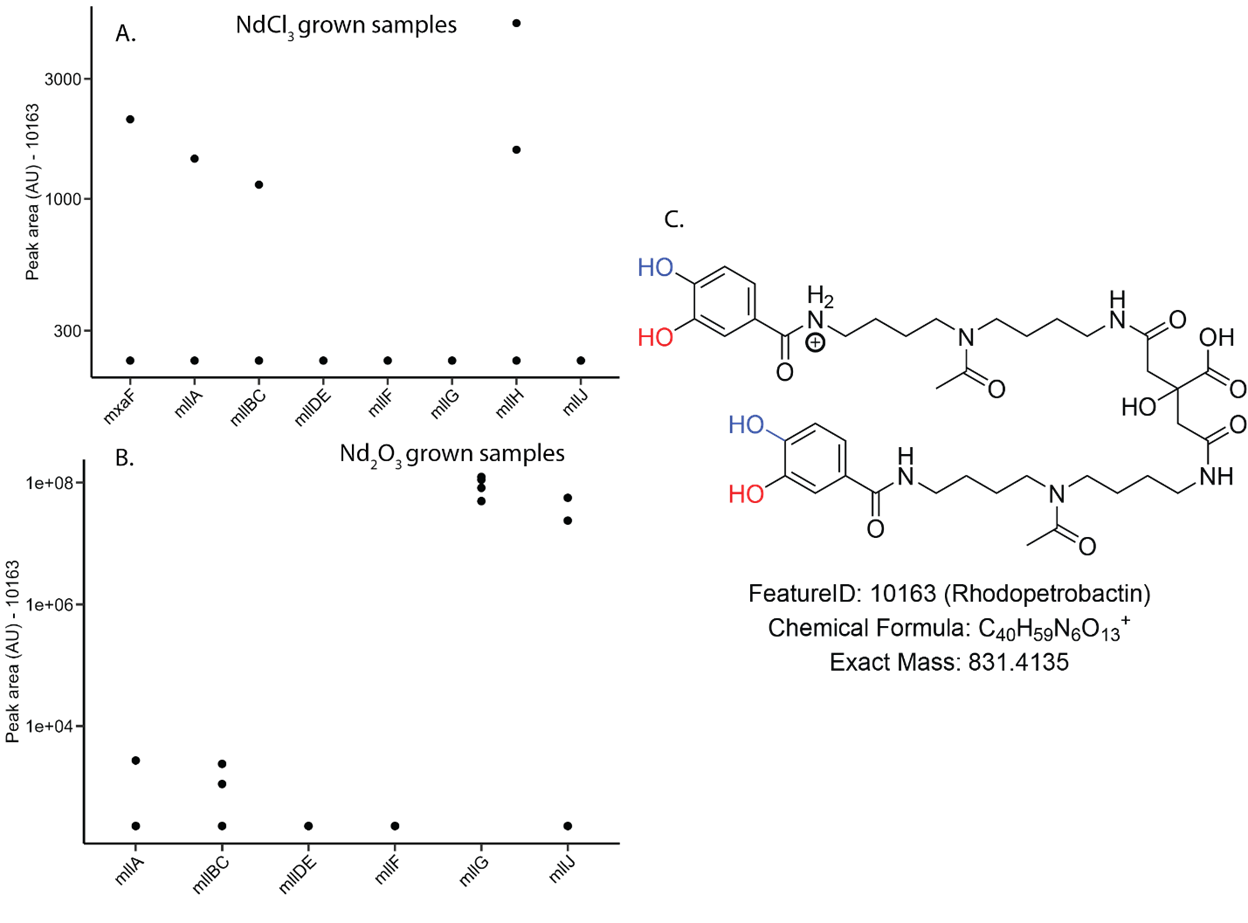

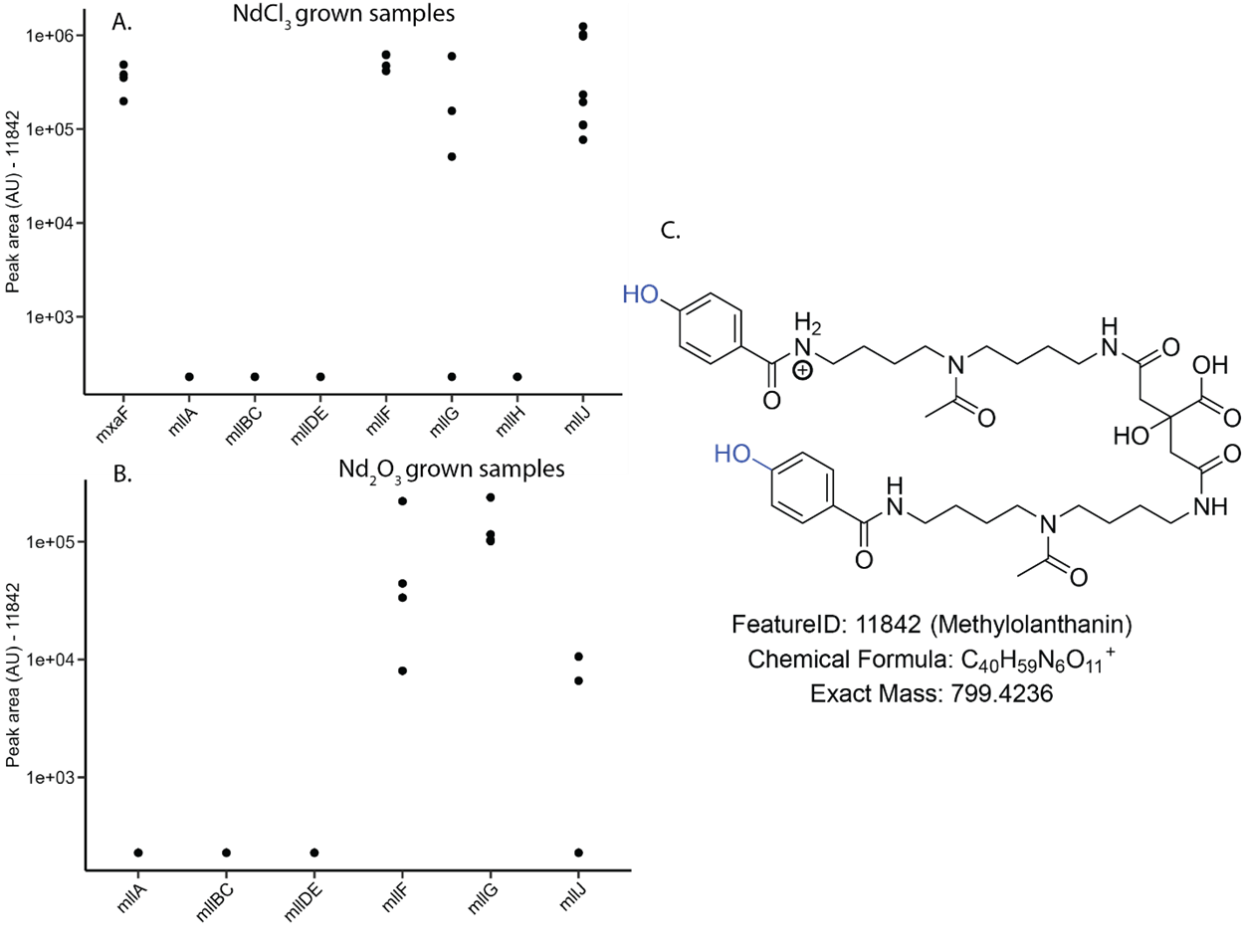

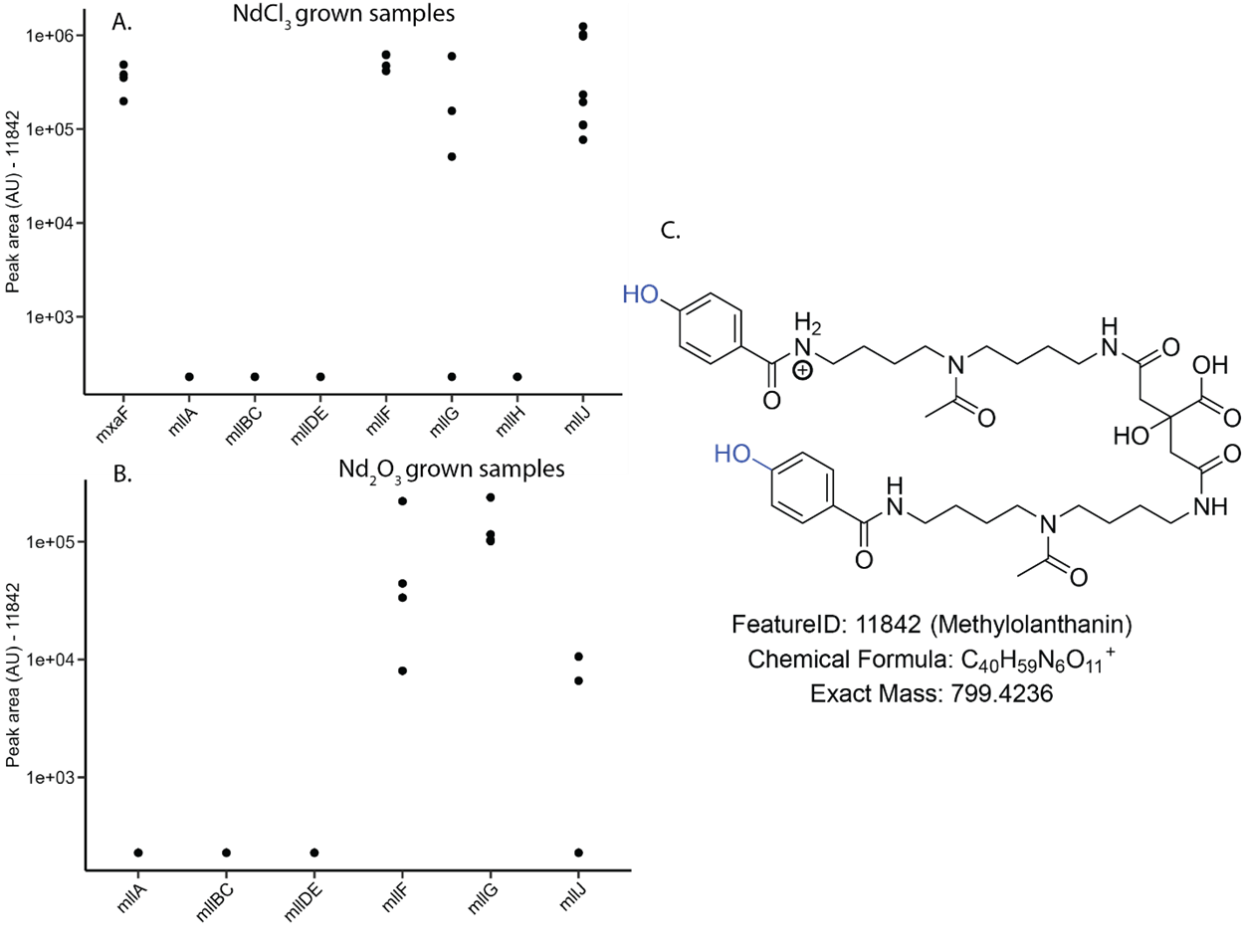

**Figure S5.** Imputed peak area plots of RPB in (A) Nd_2_O_3_ grown samples. (B) Structure of RPB (feature 10163). Low peak area for RPB in *mllA*, *mllBC,* and *mllH* mutants was nor reported as presence in these samples due to no MS/MS fragmentation spectra. This data was used to determine the presence/absence of this feature in Figure 2B. Peak areas have not been normalized to OD and originate from different sample growths.

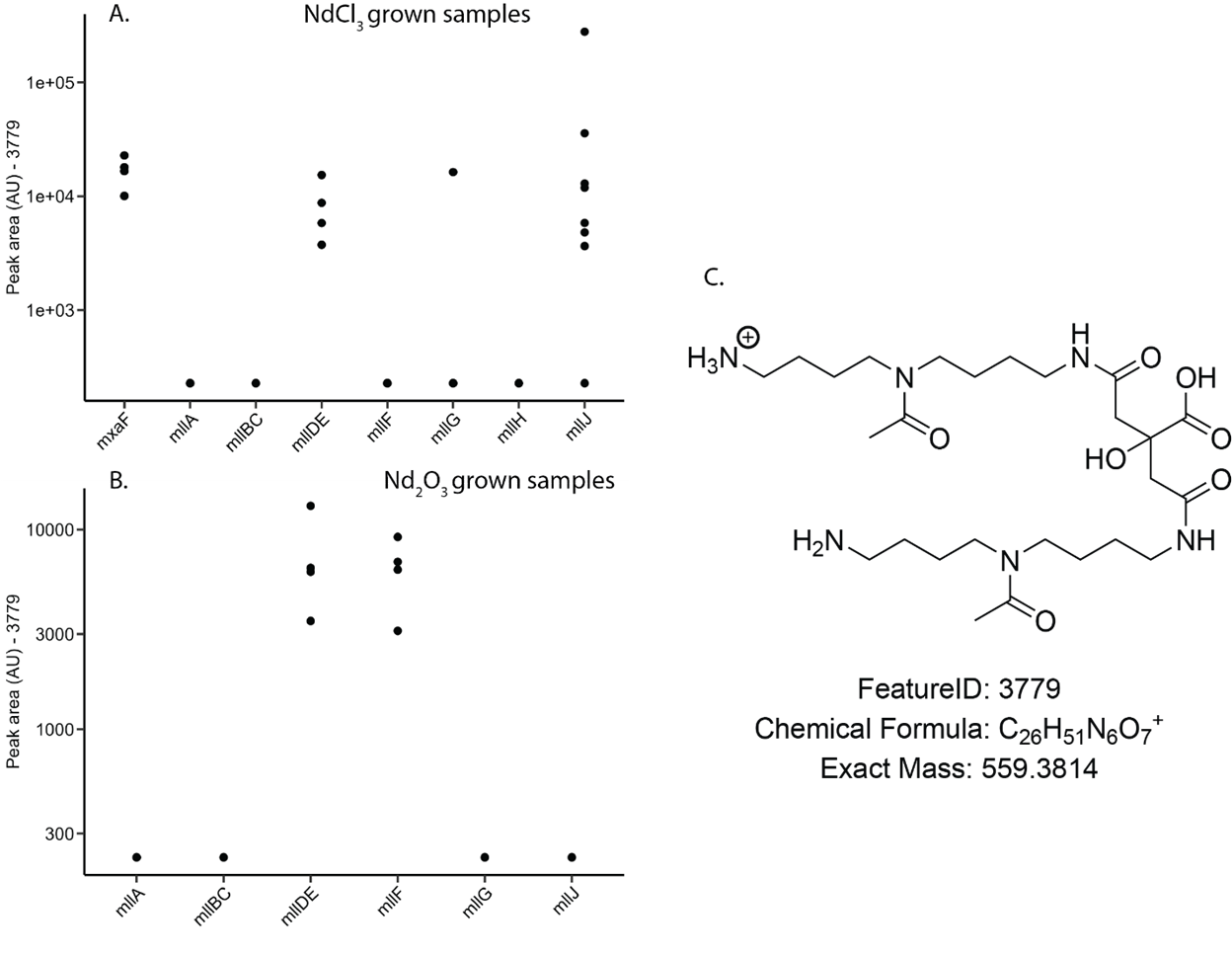

**Figure S6**. Imputed peak area plots of feature 3779 in (A) NdCl_3_ grown samples and (B) Nd_2_O_3_ grown samples. (C) Structure of feature 3779. This data was used to determine the presence/absence of this feature in Figure 2B. Peak areas have not been normalized to OD and originate from different sample growths.

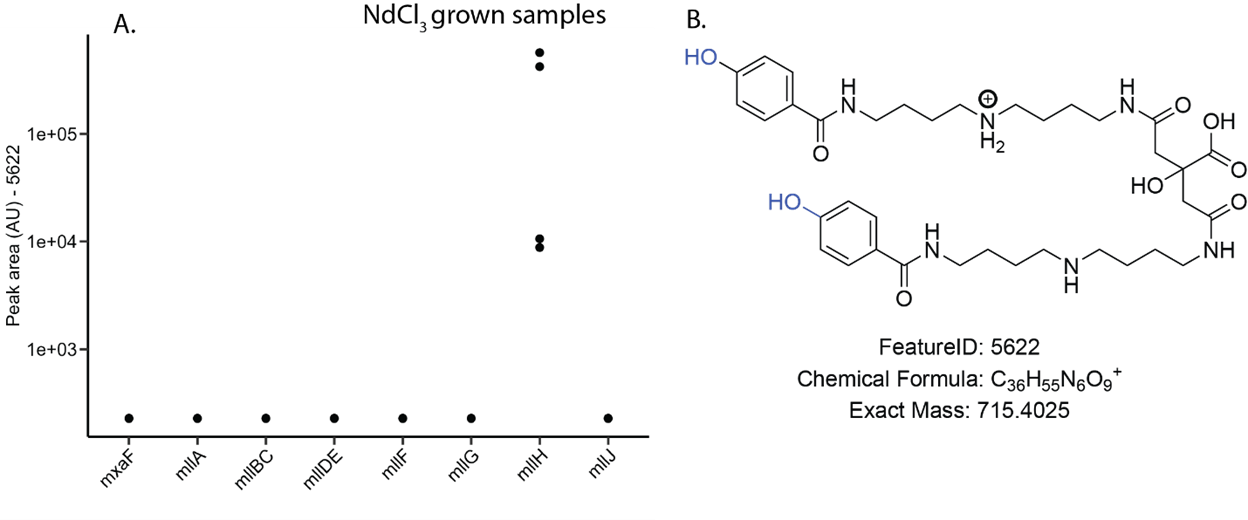

**Figure S9**. Imputed peak area plots of feature 5622 in (A) NdCl_3_ grown samples. (B) Structure of feature 5622. Peak areas have not been normalized to OD and originate from different sample growths.

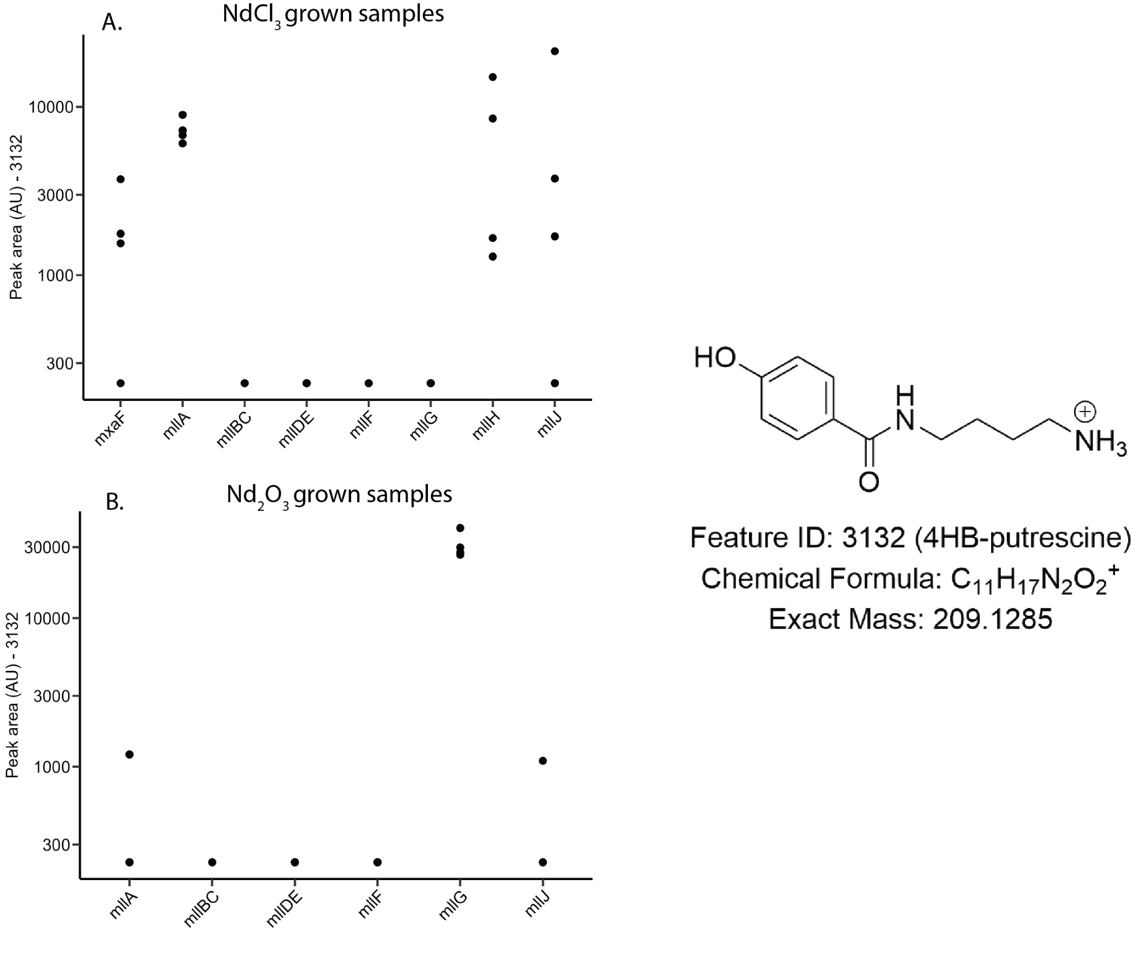

**Figure S10.** Peak area of 4-HB putrescine (feature 3132) in (A) NdCl_3_ grown samples and (B) Nd_2_O_3_ grown samples. (C) Structure of feature 3132. This data was used to determine the presence/absence of this feature in Figure 2B. Peak areas have not been normalized to OD and originate from different sample growths.

 
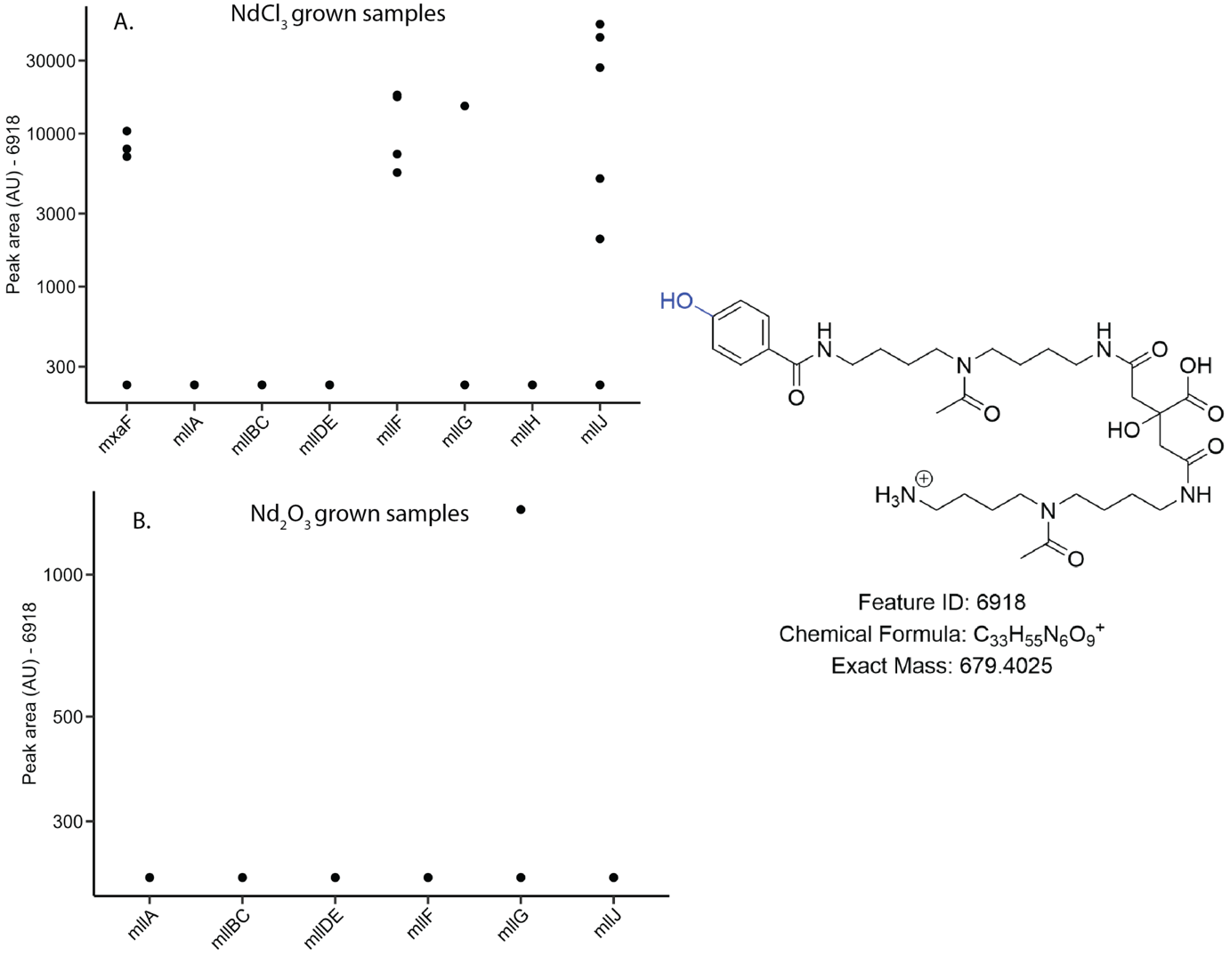

**Figure S11**. Peak area of single 4-HB methylolanthanin (feature 6918) in (A) NdCl_3_ grown samples and (B) Nd_2_O_3_ grown samples. (C) Structure of feature 6918. Peak areas have not been normalized to OD and originate from different sample growths.

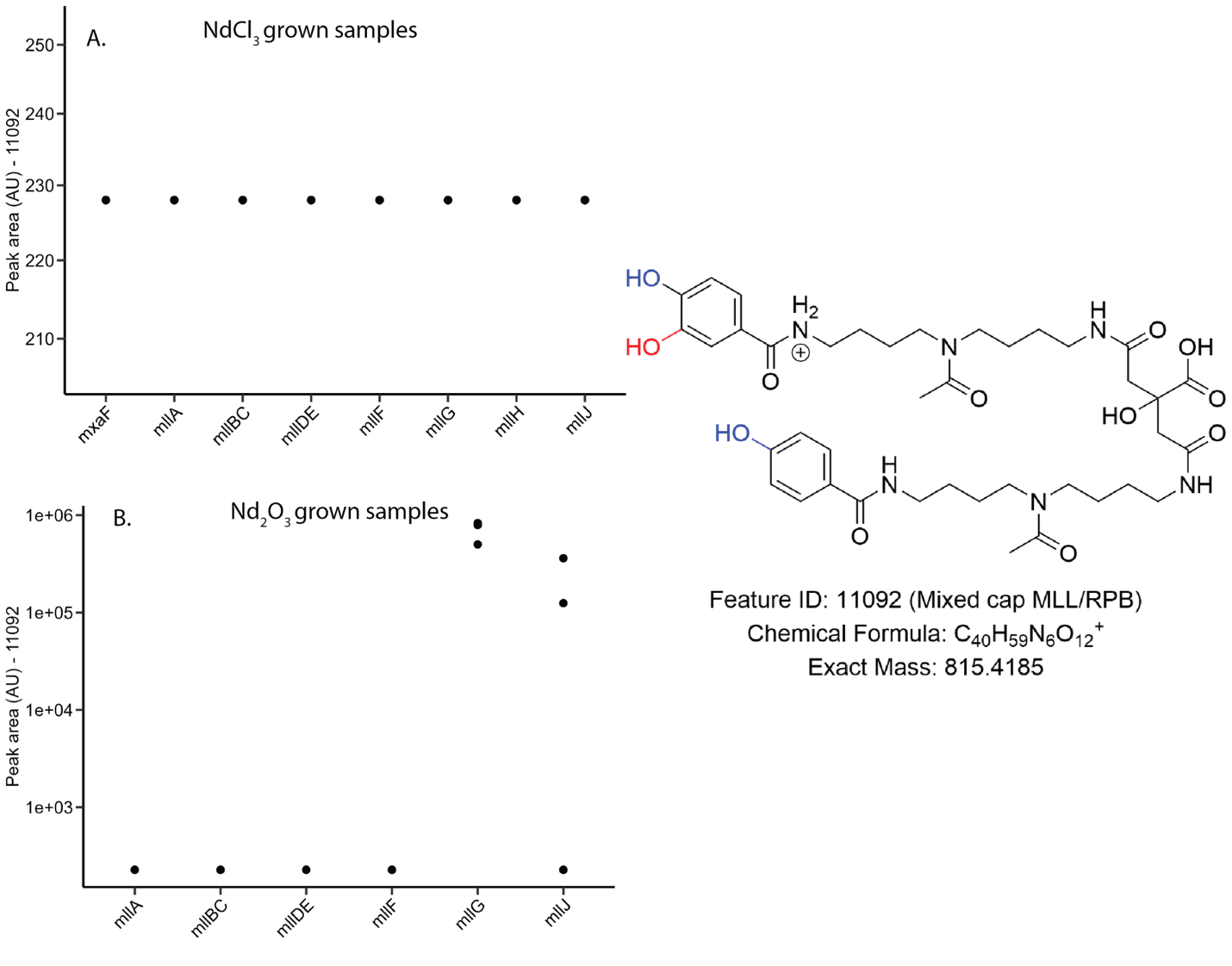

**Figure S12.** Peak area of asymmetric MLL/RPB (mixed cap chelator) in (A) NdCl_3_ grown samples and (B) Nd_2_O_3_ grown samples. (C) Structure of feature 11092. Peak areas have not been normalized to OD and originate from different sample growths.
